# Divergent roles of CPK28 in immune homeostasis across land plants

**DOI:** 10.1101/2024.10.01.616128

**Authors:** Ruoqi Dou, Baptiste Castel, Cailun A.S. Tanney, Melissa Bredow, Virginia Natali Miguel, Karima El Mahboubi, Jiashu Chu, Katharina Melkonian, Maria Camila Rodriguez Gallo, Dominique Lauressergues, Jean Keller, Márcia Gonçalves-Dias, Thomas A. DeFalco, R. Glen Uhrig, Cyril Zipfel, Pierre-Marc Delaux, Jacqueline Monaghan

## Abstract

Calcium-dependent protein kinases (CDPKs) decode cellular calcium transients and play diverse roles in plant growth and stress responses, including immunity. In *Arabidopsis thaliana* (At, Arabidopsis thereafter), AtCPK28 contributes to immune homeostasis by phosphorylating subgroup IV plant U-box proteins AtPUB22/24/25/26, which target the key immune receptor-like cytoplasmic kinase (RLCK) AtBIK1 for turnover. While this module is conserved in multiple angiosperms, it is unclear if the role of CPK28 in immune homeostasis is conserved more broadly across land plants. Here, we took an evolutionary comparative approach to understand the role of CPK28. We identified a single CPK28 ortholog in the liverwort *Marchantia polymorpha*, MpCPK28, which exhibits Ca^2+^-dependent kinase activity that is inhibited by calmodulin *in vitro*. We identified the subgroup IV plant U-box protein MpPUB20e as a substrate of MpCPK28. MpPUB20e is able to ubiquitinate MpPBLa, the functional ortholog of AtBIK1. We also provide preliminary evidence that MpPBLa undergoes proteasomal degradation in Marchantia, suggesting that optimization of MpPBLa protein accumulation is conserved across land plants. Interestingly, while loss of CPK28 function in multiple angiosperm species results in enhanced immune signaling, we find that Marchantia *Mpcpk28* mutant alleles do not display enhanced immune-triggered production of reactive oxygen species or resistance to two pathogens. However, transgenic expression of MpCPK28 was able to restore function in Arabidopsis *cpk28-1* mutants, suggesting latent functional conservation of MpCPK28. Furthermore, while AtCPK28-mediated phosphorylation of Thr95/94 on AtPUB25/26 is known to contribute to their activation, we could not observe a functional role for the equivalent residue Thr122 on MpPUB20e. Taken together, our results suggest that post-translational fine-tuning by CPK28 is likely to have refined the ‘PUB-BIK1’ module in the vascular plant lineages.

## Main text

Calcium-dependent protein kinases (CDPKs) are a unique family of Ca^2+^-regulated kinases found in apicomplexan protists (Billker *et al*., 2009), red and green algae (Valmonte *et al*., 2014; Brawley *et al*., 2017), and throughout the plant kingdom (Valmonte *et al*., 2014; Yip Delormel & Boudsocq, 2019). Thought to have evolved from the ancestral fusion between calmodulin (CaM) and CaM kinase (Chen *et al*., 2017), CDPKs contain both an N-terminal kinase domain and a C-terminal CaM-like domain bearing Ca^2+^-binding EF hands. Following divergence from a common ancestor, CDPKs in the plant kingdom separated into five phylogenetic subgroups: I, II, III, IV, and CDPK-related proteins (CRKs) that contain degenerate EF hands (Valmonte *et al*., 2014; Wang *et al*., 2016; Yip Delormel & Boudsocq, 2019) (**Figures 1A; S1**). This diversification occurred between 270-340 MYA, after terrestrial habitation and coinciding with the split between the two main land plant lineages: nonvascular and vascular plants (Valmonte *et al*., 2014). Given the diverse developmental and stress-responsive functions of CDPKs in angiosperms (Yip Delormel & Boudsocq, 2019), it is possible that expansion of the CDPK family accompanied early adaptation to terrestrial environments (Valmonte *et al*., 2014). Subgroup-IV CDPKs are the most divergent among CDPKs and were the first to diverge from the last common ancestor (Valmonte *et al*., 2014). We identified subgroup-IV CDPKs across 193 plant genomes including charophyte algae and nonvascular plants (**Figure S2; Table S1**), noting multiple lineage- and species-specific duplications that produced co-orthologs such as the Brassicaceae-specific triplication that gave rise to AtCPK16, AtCPK18, and AtCPK28. This evolutionary history suggests that lineage-specific duplications and subsequent expansion may have contributed to adaptive diversification.

**Figure 1.**
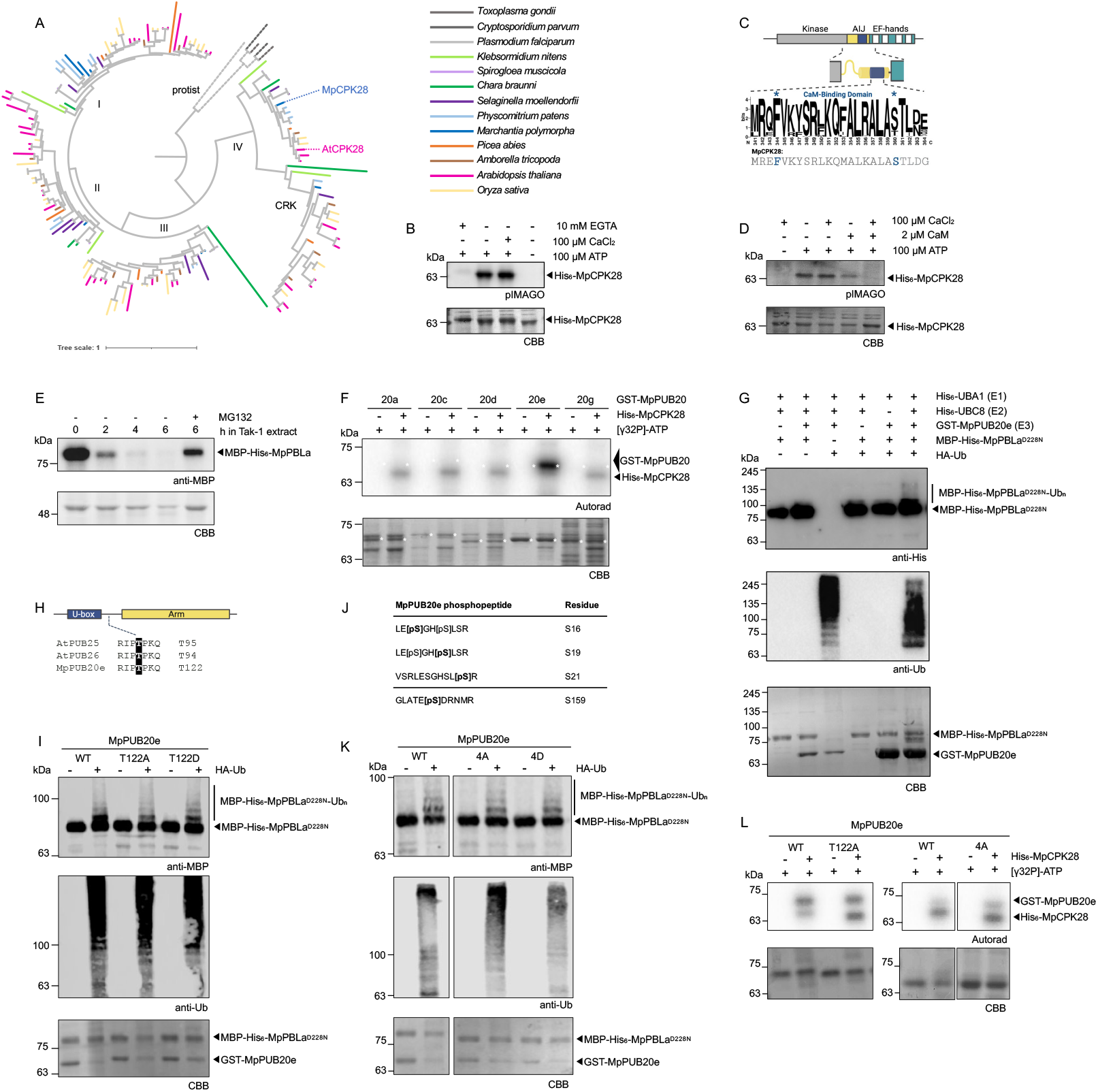
MpCPK28 is a calcium-dependent protein kinase and MpPUB20e can trans-ubiquitinate MpPBLa^D228N^. **(A)** Phylogenetic tree of CDPKs from various plant lineages, with subgroups I, II, III, IV, and CRK indicated. The tree was constructed using the Maximum Likelihood method with IQ-TREE v2.3.4 (1,000 SH-aLRT and ultraFast bootstrap replicates) and Jackknife resampling. A total of 157 CDPK protein sequences from *Selaginella moellendorffii, Physcomitrium patens, Chara braunii, Klebsormidium nitens, Spirogloea muscicola, Oryza sativa, Amborella tricopoda, Picea abies, Arabidopsis thaliana* and *Marchantia polymorpha* were aligned with MAFFT v7.490, trimmed using TrimAL v1.4.rev15, and visualized in iTOL. Five apicomplexan CDPK sequences were used as an outgroup. See **Figure S1** for a fully labeled tree. Phylogenetic analysis by VNM, visualized in iTOL. **(B)** *In vitro* kinase assays of His_6_-MpCPK28 with and without EGTA, CaCl_2_, or ATP, visualized using pIMAGO. Coomassie Brilliant Blue (CBB) indicates loading. These experiments were repeated several times with similar results by MB and RD; representative blots by MB are shown. **(C)** Schematic indicating the location of the CaM-binding domain (coloured blue) within the autoinhibitory junction (AIJ; coloured yellow) of a canonical CDPK. WebLogo indicates conservation within this region in subgroup IV CDPKs from 53 species (described in (Bredow *et al*., 2021)). The specific sequence of this region in MpCPK28 is shown below the logo. Asterisks indicate residues shown to be important for CaM binding in AtCPK28 (Bender *et al*., 2017). This figure was created in BioRender by JM. **(D)** *In vitro* kinase assays of His_6_-MpCPK28 with and without CaM, CaCl_2_, or ATP, visualized using pIMAGO. Coomassie Brilliant Blue (CBB) indicates loading. These experiments were repeated several times with similar results by RD; representative blots are shown. **(E)** Degradation of recombinant MBP-His_6_- MpPBLa following incubation for 0, 2, 4, or 6 hours in total protein extract from Marchantia Tak-1 tissue with or without the proteasome inhibitor MG132. Western blots were probed with anti-MBP antibodies, and protein loading is indicated by post-staining of Coomassie Brilliant Blue (CBB) of RuBisCO. These assays were repeated more than three times with similar results by RD. **(F)** *In vitro* kinase assays using His_6_-CPK28 as the kinase and GST-tagged MpPUB20a (‘20a’), MpPUB20c (‘20c’), MpPUB20d (‘20d’), MpPUB20e (‘20e’), or MpPUB20g (‘20g’) as substrates. Autoradiographs (Autorad) indicate incorporation of γ32P and protein loading is indicated by post-staining with Coomassie Brilliant Blue (CBB). Assays were performed more than 3 times each by RD with similar results; representative data are shown. **(G, I, K)** *In vitro* ubiquitination assays using His_6_-AtUBC1 as the E1 activating enzyme, His_6_-AtUBC8 as the E2 conjugating enzyme, GST-MpPUB20e (or indicated variants) as the E3 ligase enzyme, and MBP-His_6_-MpPBLa^D228N^ as substrate. The upper western blots are probed with anti-His or anti-MBP antibodies; the middle blot is probed with anti-Ub antibodies; and protein loading is indicated by post-staining with Coomassie Brilliant Blue (CBB). **(H)** Multiple sequence alignment of T95/T94/T122 in AtPUB25/26 or MpPUB20e. **(J)** Unique MpPUB20e phosphopeptides following kinase assays with His_6_-CPK28. Each peptide was present in at least 2/3 independent replicates and not found in control samples with ATP but without His_6_-CPK28. The bolded sites are the phosphosites and their positions are indicated on the right. Kinase assays performed by RD; trypsin digests and LC-MS/MS performed by MCRG. **(L)** *In vitro* kinase assays using His_6_-CPK28 as the kinase and GST-tagged MpPUB20e and mutant variants as substrates. Autoradiographs (Autorad) indicate incorporation of γ32P and protein loading is indicated by post-staining with Coomassie Brilliant Blue (CBB). Assays were performed more than 3 times each by RD with similar results; representative data are shown. Cloning credits are provided in **Table S1**.

CPK28 regulates immune homeostasis and growth in multiple angiosperms including Arabidopsis (Matschi *et al*., 2013, 2015; Monaghan *et al*., 2014; Jin *et al*., 2017; Dressano *et al*., 2020; Bredow *et al*., 2021; Liu *et al*., 2022; Ding *et al*., 2022b; Sowders *et al*., 2024), tomato (Hu *et al*., 2021; Ding *et al*., 2022a), rice (Campos-Soriano *et al*., 2011; Bundó & Coca, 2016; Wang *et al*., 2018b; Li *et al*., 2022), and cotton (Wu *et al*., 2021; Gao *et al*., 2024; Wang *et al*., 2024). We took advantage of recent advances in functional genomics in *Marchantia polymorpha* (hereafter, Marchantia) to test the role of CPK28 in a nonvascular plant. We identified a single copy of *MpCPK28* in Marchantia (**Figures 1A; S1; S2; S3A**), that is broadly expressed in both the thallus and sexual reproductive tissues (**Figure S3B**). MpCPK28 is an active kinase with a low Ca^2+^ threshold for activation, as its autophosphorylation activity is only marginally increased with the addition of CaCl_2_ but is strongly reduced in the presence of the Ca^2+^ chelator EGTA (**Figure 1B**) – mirroring AtCPK28 (Matschi *et al*., 2013; Bender *et al*., 2017; Bredow *et al*., 2021). Among CDPKs, AtCPK28 is uniquely inhibited by Ca^2+^-dependent interactions with CaM (Bender *et al*., 2017). Residues critical for this interaction, Phe344 and Ser360, are highly conserved across plants, including MpCPK28 (**Figure 1C**). Consistent with this, CaM inhibited MpCPK28 autophosphorylation (**Figure 1D**), suggesting this inhibitory mechanism is an ancient property of subgroup-IV CDPKs.

In Arabidopsis, AtCPK28 phosphorylates the E3 ubiquitin ligases AtPUB22, AtPUB24, AtPUB25 and AtPUB26 (Wang *et al*., 2018a; Dou *et al*., 2025). This contributes to their polyubiquitination of AtBIK1, which is degraded by the 26S proteasome to optimize immune signaling (Monaghan *et al*., 2014; Wang *et al*., 2018a). As MpPBLa is the functional ortholog of AtBIK1 (Chu *et al*., 2023), we tested if it similarly undergoes proteasomal degradation. We found that both MpPBLa and AtBIK1, but not another RLCK protein, AtCARK7 (Gonçalves Dias *et al*., 2025), were rapidly degraded in cell-free Marchantia Tak-1 protein extracts within 2 hours and stabilized by the addition of proteasome inhibitor MG132 (**Figures 1E; S4A-B**). This suggests that Marchantia contains conserved machinery capable of degrading MpPBLa and AtBIK1.

We hypothesized that this machinery may include Marchantia orthologs of AtPUB22/24/25/26. A recent phylogeny of the PUB family across land plants (Trenner *et al*., 2022) indicates that, although none of the 38 MpPUBs form a monophyletic group with AtPUB25/26 or related AtPUB22/23/24, seven are monophyletic with the sister group AtPUB20/21, which we name MpPUB20a-g (sharing 30-70% amino acid sequence identity) (**Figure S5A**). Based on their phylogenetic relationship and gene expression profiles (**Figure S5B**), we chose five to test as MpCPK28 substrates: MpPUB20a, MpPUB20c, MpPUB20d, MpPUB20e, and MpPUB20g. Although AtCPK28 phosphorylates multiple PUBs (Wang *et al*., 2018a; Dou *et al*., 2025), we found that MpCPK28 was only able to phosphorylate MpPUB20e (**Figure 1F**). This may be due to the high divergence between MpPUB20 proteins, which share 29-69% sequence identity.

Previous work showed that AtPUB22 interacts with multiple subgroup-IV E2 ubiquitin conjugating enzymes (UBCs) (Turek *et al*., 2018), and that AtPUB22/24 and AtPUB25/26 can pair with AtUBC8 (Wang *et al*., 2018a; Dou *et al*., 2025). We found that MpPUB20e can autoubiquitinate when paired with either AtUBC8 or AtUBC30 (sharing 96% and 85% amino acid sequence identity to Marchantia UBC protein Mp5g21800, respectively) (**Figure S6**). Whilst we were unable to detect clear ubiquitination by MpPUB20e on MpPBLa (**Figure S7**), we were able to detect ubiquitination on the catalytically inactive mutant MpPBLa^D228N^ (**Figure 1G**). This is notable since AtPUB22/24/25/26 preferentially ubiquitinate inactive AtBIK1 in Arabidopsis (Wang *et al*., 2018a; Dou *et al*., 2025), and suggests that MpPUB20e also preferentially targets inactive MpPBLa. These results suggest that MpPBLa is under proteasomal control in Marchantia and can be ubiquitinated by MpPUB20e.

In Arabidopsis, AtCPK28 phosphorylates AtPUB25/26 on Thr95/94 which partially contributes to their activity (Wang *et al*., 2018a). Although residue T122 in MpPUB20e is in a conserved position (**Figure 1H**), we did not detect any impact on ubiquitination of MpPBLa^D228N^ using either phospho-ablative MpPUB20e^T122A^ or phospho-mimetic MpPUB20e^T122D^ variants (**Figure 1I**). Using mass spectrometry, we identified four MpCPK28-mediated phosphosites on MpPUB20e: Ser16, Ser19, and Ser21 at the far N-terminus, and Ser159 at the start of the Armadillo repeat domain (**Figure 1J**). We generated phospho-ablative MpPUB20e^4A^ and phospho-mimetic MpPUB20e^4D^ variants (representing all four sites we identified mutated to Ala or Asp) but again did not observe any differences in ubiquitination of MpPBLa^D228N^ (**Figure 1K**). Although this suggests that MpCPK28-mediated phosphorylation of MpPUB20e may not impact its function, it is important to note that MpCPK28 phosphorylates MpPUB20e^4A^ and MpPUB20e^T122A^ similarly to MpPUB20e (**Figure 1L**), indicating that we did not identify all phosphosites.

In any case, as we did not observe any contribution of T122 to MpPUB20e function, we propose that post-translational fine-tuning of AtPUB25/26 at T95/94 may have evolved after the divergence between nonvascular and vascular plants.

To assess the function of MpCPK28 in immune signaling, we isolated three CRISPR-generated null mutants: *Mpcpk28-11, Mpcpk28-14*, and *Mpcpk28-21* (**Figure 2A**), all of which grew smaller than wild-type Tak-1 (**Figure 2B,C**). We also attempted to generate *MpCPK28* overexpression lines, however we were only able to isolate viable gemmae from one line, MpCPK28-OE2, in which *MpCPK28* is expressed ∼30 times higher than in wild-type Tak-1 (**Figure S8A**). MpCPK28-OE2 displayed defects in both gemmae (**Figure S8B**) and thallus (**Figure S8F**) morphology, containing additional and irregularly distributed apical notches (**Figure S8C-D**). This phenotype is reminiscent of dysfunctional hormone signaling (Liang *et al*., 2022; Takahashi *et al*., 2026). Although we could not analyze additional lines, these data suggest that genetic manipulation of *MpCPK28* disrupts growth and/or developmental programs. This is notable, as genetic manipulation of CPK28 orthologs in multiple angiosperm species also results in developmental defects. Arabidopsis loss-of-function *cpk28* mutants are defective in the transition from vegetative-to-reproductive growth, presenting with delayed and stunted stem elongation (Matschi *et al*., 2013). Similarly, loss-of-function *cpk28* mutants in tomato (Hu *et al*., 2021) and rice (Li *et al*., 2022) also display smaller growth. Together, these data suggest a conserved role for CPK28 in growth and development across land plants.

**Figure 2.**
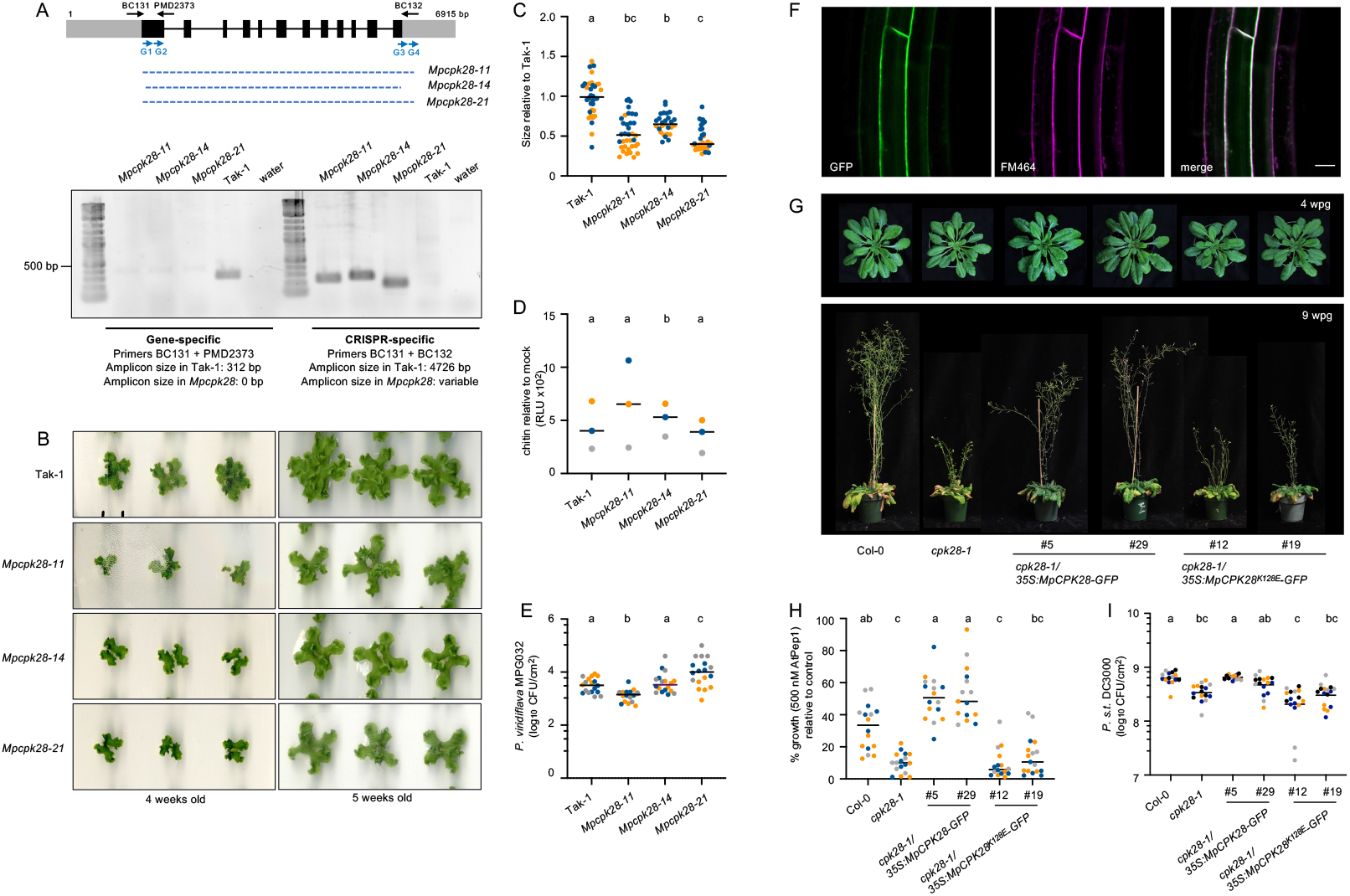
Functional analysis of *MpCPK28* in Marchantia and Arabidopsis. **(A)** Schematic representation of the MpCPK28 locus with exons and introns indicated by boxes and lines, respectively. Grey regions on the termini indicate 5’ and 3’ untranslated regions. The location and orientation of guide RNAs (G1, G2, G3, G4) used in the generation of CRISPR lines are indicated with blue arrows below the gene schematic, and the area deleted in *Mpcpk28-11, Mpcpk28-14*, and *Mpcpk28-21* mutant alleles is indicated by a dashed blue line. DNA gel electrophoresis of amplicons resulting from polymerase chain reactions using primers indicated above the gene schematic. The 500 bp molecular marker is indicated; sequences for primers and cloning credits are provided in **Table S1. (B-C)** Photographs **(B)** and quantification of thallus size **(C)** of the *Mpcpk28* mutant alleles compared to Tak-1 grown on agar plates. Values in C are the thallus sizes of individuals relative to Tak-1 at 4 weeks old; the different colours represent independent experiments. Statistically significant groups are indicated with lower-case letters following a one-way analysis of variance (ANOVA) followed by Tukey’s post-hoc test (p<0.001). Experiments conducted by BS. **(D)** ROS production measured in relative light units (RLUs) after treatment with 1 µM 7-mer chitin. Values are mean RLU relative to mock (n=3, with 5 gemmalings pooled per sample), colour-coded by experimental replicate; the line represents the mean of all experiments together. Statistically significant groups are indicated with lower-case letters following a one-way analysis of variance (ANOVA) followed by Tukey’s post-hoc test (*p*<0.01). Experiments conducted twice by KM and once by RD. **(E)** Growth of *Pseudomonas viridiflava* MPG032 in the indicated genotypes at 3 days post inoculation. Values are individual data points colour-coded by experimental replicate (log_10_ colony forming units per cm^2^ area; n=6 thallus discs per experiment) and the line represents the mean of all experiments together. Statistically significant groups are indicated with lower-case letters following a one-way analysis of variance (ANOVA) followed by Tukey’s post-hoc test (*p*<0.01). Experiments performed by BS. **(F)** Single-plane confocal micrographs of root cells from *cpk28-1/35S:MpCPK28-GFP* line #5 indicating the subcellular localization of MpCPK28-GFP (GFP) and the plasma membrane (FM464). Scale bar indicates 10 μm. Data collected by RD. **(G)** Photos of Arabidopsis plants at 4 (upper) and 9 (lower) weeks post germination (wpg) grown under short-day conditions. Similar observations were made across generations over multiple years by RD, CAST, and MB; photos taken by RD. **(H)** Seedling growth inhibition assay indicating % growth in media containing 500 nM AtPep1 peptide relative to growth in media without peptide. Values represent % fresh weight of single seedlings (n=5-6 for each treatment per genotype) colour-coded by experimental rep. Statistically significant groups were evaluated by a one-way ANOVA followed by a Tukey’s post-hoc test and are indicated by lower-case letters (*p*<0.0001). Data collected by VNM. **(I)** Growth of *Pseudomonas syringae* pv. *tomato* (*P*.*s*.*t*.) isolate DC3000 3 days after syringe-inoculation, colour-coded by experiment. Values are colony forming units (cfu) per leaf area (cm^2^) from 4-5 samples per genotype (each sample contains 3 leaf discs from 3 different infected plants). The line represents the mean. Lower case letters indicate that all genotypes are in the same statistical group, as determined by a one-way ANOVA followed by Tukey’s post-hoc test (p<0.002). Data collected by VNM. Details regarding plant genotypes are provided in **Table S1**.

Loss-of-function *cpk28* mutants in Arabidopsis (Monaghan *et al*., 2014; Sowders *et al*., 2024), rice (Wang *et al*., 2018b), cotton (Wu *et al*., 2021; Wang *et al*., 2024), and tomato (Ding *et al*., 2022a), display enhanced immune signaling and anti-microbial resistance, while overexpression of *AtCPK28* in Arabidopsis strongly reduces immune responses (Monaghan *et al*., 2014). A conserved hallmark of immune induction in plants is a rapid burst of apoplastic reactive oxygen species (ROS) catalyzed by NADPH oxidases (DeFalco & Zipfel, 2021; Chu *et al*., 2023; Yotsui *et al*., 2023). Functionally orthologous to AtBIK1 (Kadota *et al*., 2014), MpPBLa directly phosphorylates conserved N-terminal residues on the NADPH oxidase MpRBOH1 required for its function (Chu *et al*., 2023). We observed slightly higher chitin-induced ROS levels in *Mpcpk28-14* compared to wild-type Tak-1 over three independent trials but did not observe any differences in *Mpcpk28-11* or *Mpcpk28-21* (**Figure 2D**), suggesting that loss of *MpCPK28* does not measurably impact chitin-induced ROS. Comparatively, we consistently observed lower chitin-induced ROS in MpCPK28-OE2 compared to wild-type Tak-1 across multiple independent biological experiments (**Figure S8E-F**). Unfortunately, the MpCPK28-OE2 line lost viability over time and we were unable to generate additional lines. However, we were able to infect the *Mpcpk28* mutants with the bacterial pathogen *Pseudomonas viridiflava* isolate MPG032 (Robinson *et al*., 2025). While we observed lower bacterial growth after 3 days in *Mpcpk28-11*, we observed higher growth in *Mpcpk28-21* and no difference in *Mpcpk28-14* compared to wild-type Tak-1 (**Figure 2E**). We also infected the mutants with the fungal pathogen *Collectotrichum nymphaeae* (El Mahboubi *et al*., 2026), but were unable to obtain reproducible results across multiple independent biological experiments (**Figure S9**). As all three lines contain large deletions of *MpCPK28*, this variability is likely to reflect other genetic differences between the lines, such as the insertion loci of the CRISPR/Cas9 module. These results suggest that *MpCPK28* may not play a critical role in the Marchantia immune system (at least for the bioassays and pathogens tested), which in turn suggests that a role in immune homeostasis is a derived feature of CPK28 in vascular plants. We note that all three *Mpcpk28* null mutants did consistently grow smaller compared to wild-type Tak-1 (**Figure 2B-C**), providing robust evidence that a role in growth and development is conserved between Marchantia and angiosperms. Although speculative, this could indicate that the ancestral function of subgroup-IV CDPKs is in growth regulation and that a role in immune homeostasis arose specifically in vascular plants.

To assess if MpCPK28 could function in place of AtCPK28, we transformed Arabidopsis *cpk28-1* mutants with MpCPK28 C-terminally tagged with green fluorescent protein (GFP). We additionally transformed *cpk28-1* plants with the mutant variant MpCPK28^K128E^ that is predicted to be catalytically inactive due to the loss of a critical lysine in the ATP-binding site. We isolated homozygous transgenic lines and confirmed protein accumulation by immunoblot (**Figure S10**). MpCPK28-GFP and MpCPK28^K128E^-GFP localize to the plasma membrane (**Figures 2F, S11**), similar to AtCPK28 (Matschi *et al*., 2013). Arabidopsis mutant *cpk28-1* plants display both enhanced immune signaling (Monaghan *et al*., 2014) as well as defects in stem elongation (Matschi *et al*., 2013). While MpCPK28-GFP complemented the stem elongation defect in *cpk28-1*, the catalytically inactive variant did not (**Figure 2G**), suggesting that MpCPK28 can replace AtCPK28 during the vegetative-to-reproductive stage transition. Likewise, MpCPK28, but not MpCPK28^K128E^, complemented enhanced immune-triggered seedling growth inhibition (**Figure 2H**), and resistance to the bacterial pathogen *Pseudomonas syringae* pv. tomato DC3000 (**Figure 2I**) in *cpk28-1*. These results suggest that MpCPK28 can function equivalently to AtCPK28 in Arabidopsis.

We conclude that while the biochemical function of CPK28 is conserved across land plants, the post-translational fine-tuning of the PUB-BIK1/PBL module provided by CPK28 is likely to be a derived feature of this module. Although the loss of *MpCPK28* did not have a measurable impact on pathogen resistance in Marchantia, MpCPK28 is fully functional in Arabidopsis, consistent with latent functional conservation. It is possible that CPK28-mediated phosphorylation of T95/94 in AtPUB25/26 became functionally relevant as the module evolved. In Arabidopsis, phosphorylation of T95/94 is not a prerequisite for, but rather contributes to, AtPUB25/26 activation and subsequent ubiquitination and proteasomal turnover of AtBIK1 (Wang *et al*., 2018a) – suggestive of a fine-tuning mechanism. Mutation of T122 did not have a measurable impact on the ability of MpPUB20e to ubiquitinate MpPBLa^D228N^, indicating that this level of fine-tuning evolved later. More work is needed to understand how post-translational modifications evolved to regulate this critical immune module in angiosperms.

## Acknowledgements

We thank all members of the Monaghan Lab for their commitment to fostering a welcoming and collaborative research environment. Queen’s University is situated on the territory of the Haudenosaunee and Anishinaabek and we are grateful to live, work, and play on these lands. We are grateful to three anonymous peer reviewers who critically evaluated an earlier version of this manuscript and provided thoughtful suggestions to improve it. We thank Dale Kristensen and Saeid Mobini for managing the Queen’s University Phytotron Facility; Tony Papanicolaou for managing the Queen’s University Department of Biology Microscopy Suite; Jack Moore for technical assistance and maintenance of the University of Alberta Mass Spectrometry and Proteomics Facility; Anamika Rawat for training assistance in confocal microscopy; Brandon Saltzman for preliminary work on the MpPUB phylogeny; Katherine Dunning for establishing the cell-free degradation assay in the Monaghan Lab; Sarah Ruiz for assistance measuring seedling growth inhibition; and Kathy Sihavong for assisting with cloning MpCPK28. We thank Phil Carella for providing the *P. viridiflava* strain and the Genotoul Bioinformatics Platform Toulouse Occitanie (Bioinfo Genotoul, https://doi.org/10.15454/1.5572369328961167E12) for providing computing resources. This work was funded by grants awarded to **JM**: Canadian Natural Sciences and Engineering Research Council of Canada (NSERC) Discovery and Discovery Accelerator Programs (RGPIN-2016-04787, RGPAS-492902-2016, and RGPIN-2024-04072), the Canada Research Chair (CRC) Program, and Queen’s University; as well as grants awarded to **PMD**: Labex TULIP (ANR-10-LABX-41), the ‘École Universitaire de Recherche (EUR)’ TULIP-GS (ANR-18-EURE-0019), the European Research Council (ERC) under the European Union’s Horizon 2020 research and innovation programme (grant agreement no. 101001675 - ORIGINS), the project Enabling Nutrient Symbioses in Africa (ENSA), that is funded by Bill & Melinda Gates Agricultural Innovations (INV-57461), the Bill & Melinda Gates Foundation and the Foreign, Commonwealth and Development Office (INV-55767); grants to **CZ**: Swiss National Science Foundation (grant agreement no. 31003A_182625), Gatsby Charitable Foundation, and the University of Zurich; and grants to **RGU**: the Canadian Foundation for Innovation (CFI) John R. Evans Leaders Fund (JELF; grants 41831 and 37833). **MB** was supported by an NSERC Postdoctoral Fellowship (2019-2021). **TAD** was supported by a Long-Term Fellowship from the European Molecular Biology Organization (EMBO LTF no. 100-2017; 2017-2019) and an NSERC Postdoctoral Fellowship (2019-2021). **MCRG** was supported by an Alberta Graduate Excellence Scholarship (AGES). **KM** was supported by the German Research Foundation (DFG Walter Benjamin fellowship project number 536856410).

## Author Contributions

**JM** and **PMD** conceived and designed the project. **RD, BC, CAST, MB, VNM, KEM, JC, KM, MCRG, DL, MGD, JM** generated materials, performed experiments, and/or analyzed data. **JK** and **VNM** performed phylogenetics. **JM, PMD, RGU, TAD, CZ** supervised the project. **JM** wrote the manuscript with input from all authors.

## Data Availability

Any materials described in this article will be made freely available upon request. The person responsible for sharing materials is the author of correspondence. Proteomics data has been deposited to the ProteomeXchange Consortium via the PRIDE (Perez-Riverol et al., 2019) partner repository.

