## Supporting Information for "Divergent roles of CPK28 in immune homeostasis across land plants"

**The following Supporting Information is available for this article:**

- **Figure S1.** Phylogenetic tree of CDPKs from various plant lineages.
- **Figure S2.** Phylogenetic tree of subgroup IV CDPKs across the plant lineage.
- **Figure S3.** Genomic and gene expression data for MpCPK28.
- **Figure S4.** Cell-free degradation assay of AtBIK1 and AtCARK7 in Tak-1 protein extract.
- **Figure S5.** Identification of MpPUB20 genes in *Marchantia polymorpha*.
- **Figure S6.** MpPUB20e is capable of auto-ubiquitination *in vitro* when paired with E2s AtUBC8 or AtUBC30.
- **Figure S7.** *In vitro* trans-ubiquitination of MpPBLa by MpPUB20e is not detectable.
- **Figure S8**. Characterization of MpCPK28-OE2 overexpression line in *Marchantia polymorpha*.
- **Figure S9.**
- **Figure S10.** Western blot indicating MpCPK28-GFP expression in Arabidopsis transgenic lines.
- **Figure S11**. Subcellular localization of MpCPK28$^{K128E}$-GFP.

- **Methods S1**

- **Table S1.** Germplasm, clones, primers, and resources used in this study.

  *Provided as a separate spreadsheet.*

- **Supplemental References**

### Supporting Information - Figures

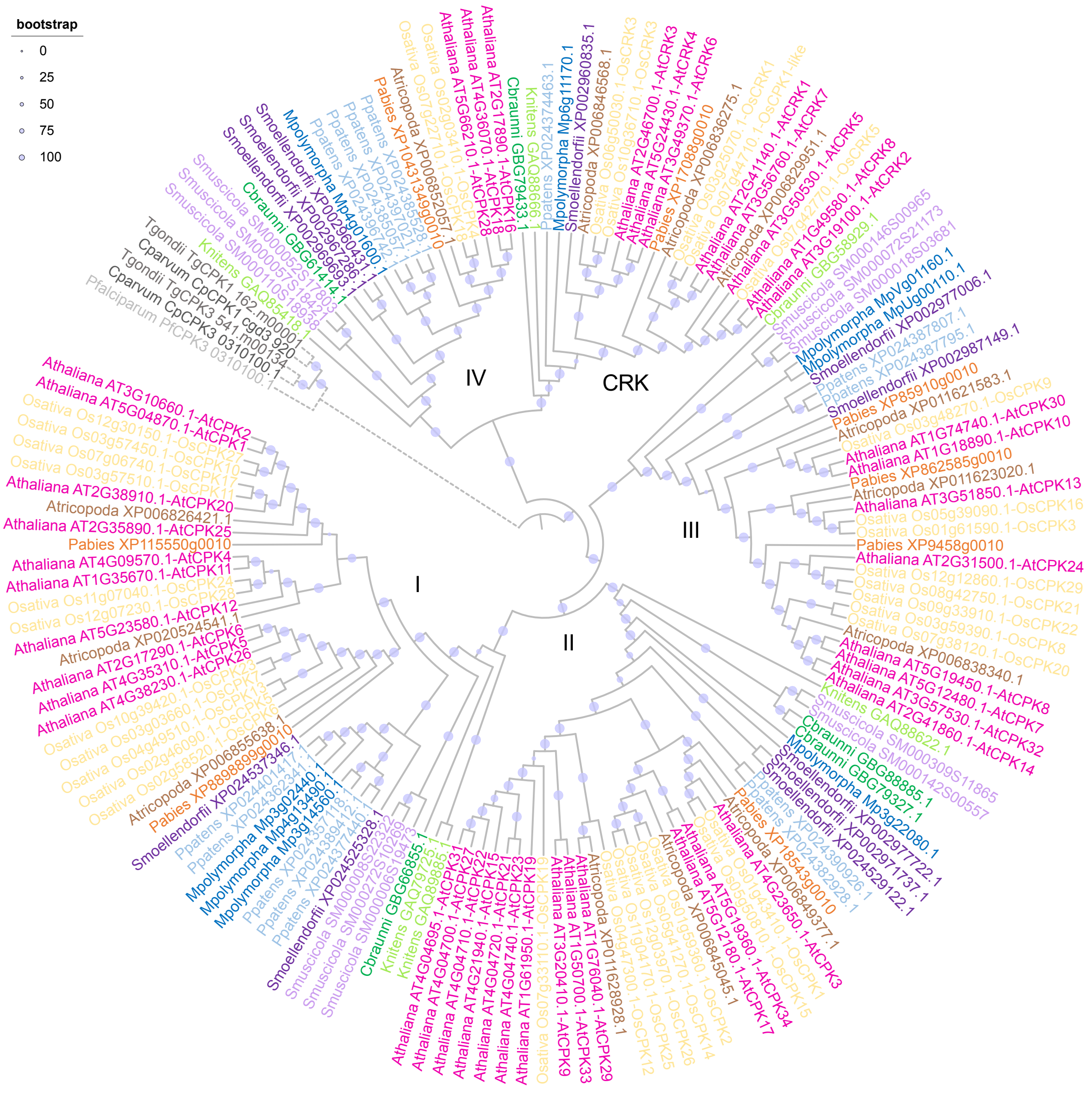


**Figure S1.** Phylogenetic tree of CDPKs from various plant lineages.

A more complete view of the phylogenetic tree presented in **Figure 1A**. CDPKs from subgroups I, II, III, IV, and CRK from *Selaginella moellendorffii, Physcomitrium patens, Chara braunii, Klebsormidium nitens, Spirogloea muscicola, Oryza sativa, Amborella tricopoda, Picea abies, Arabidopsis thaliana* and *Marchantia polymorpha*, as well as five apicomplexan CDPK sequences used as an outgroup. Gray circles at the nodes represent support from 1,000 bootstraps. Tree generated by VNM and visualized in iTOL.

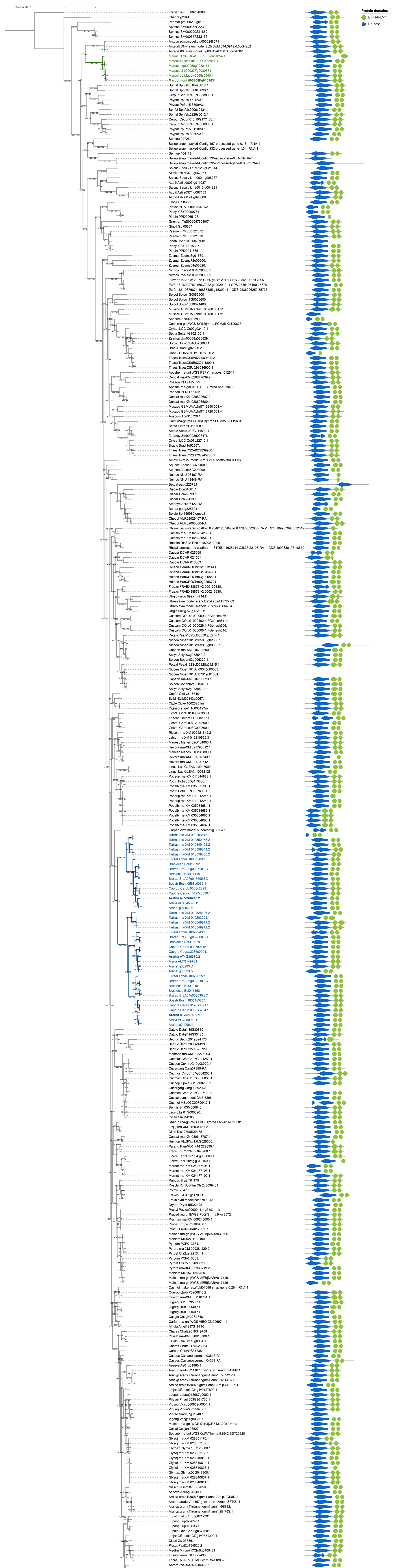


**Figure S2.** Phylogenetic tree of subgroup IV CDPKs across the plant lineage.

Maximum likelihood tree rooted on the charophyte algae Klebsormidium nitens (model: GTR+F+R7, log-likelihood: -146591.9493). Node supports are shown on the branches (sh-aLRT/UltraFast Bootstraps). Functional domains were predicted using HMMER and are depicted on the right side of the tree. EF-HAND-7 (PF13499) and Protein Kinase (PF00069) domains are indicated by green octagons and blue left-pointing pentagrams respectively. Domains are proportional to protein length and protein backbone indicated by gray line. The Marchantia clade is coloured in green, and the Brassicaceae clade is coloured in blue; *Marchantia polymorpha* subsp. *ruderalis* and *Arabidopsis thaliana* CPK28 are indicated in bold. This tree was generated by JK and visualized in iTOL.

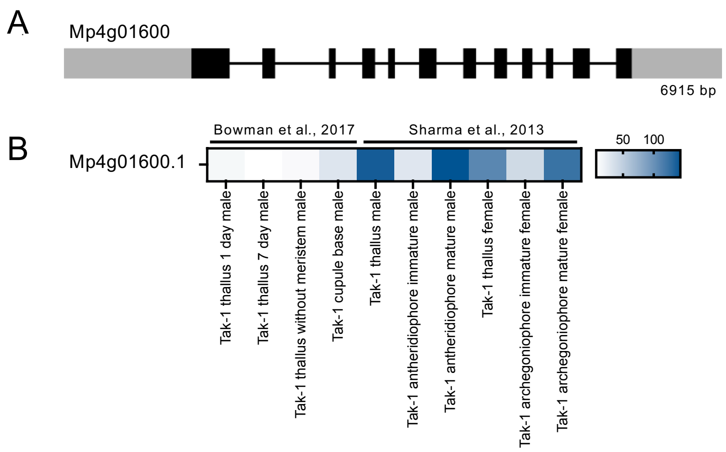


**Figure S3.** Genomic and gene expression data for *MpCPK28*.

**(A)** Schematic representation drawn to scale of *Marchantia polymorpha* gene MpCPK28/Mp4g01600.1 indicating exons (black boxes), 5' and 3' untranslated regions (gray boxes), and introns (lines). **(B)** Heatmap showing expression data of MpCPK28/Mp4g01600.1 in the indicated tissues, based on data collected from two independent studies (Sharma *et al.*, 2013; Bowman *et al.*, 2022). Data was retrieved from MarpolBase and analyzed by JM.

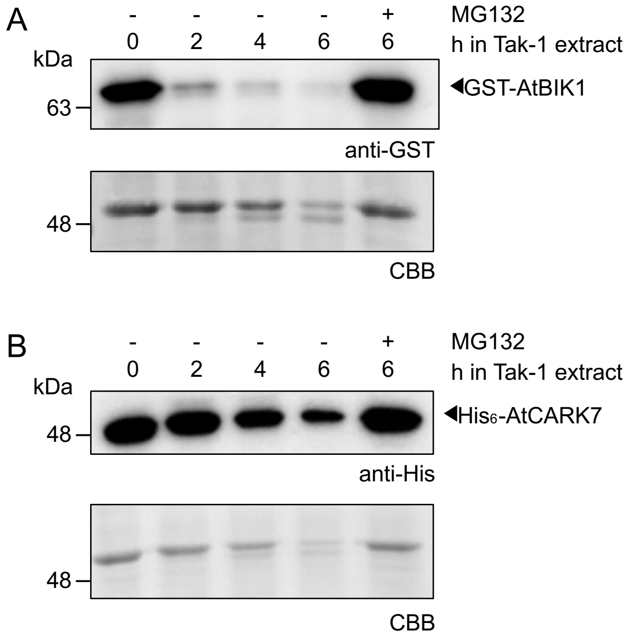


**Figure S4.** Cell-free degradation assay of AtBIK1 and AtCARK7 in Tak-1 protein extract.

Degradation of recombinant GST-AtBIK1 **(A)** or $His_6$-AtCARK7 (**B)** following incubation for 0, 2, 4, or 6 hours in total protein extract from Marchantia Tak-1 tissue with or without the proteasome inhibitor MG132. Western blots were probed with either anti-MBP or anti-GST antibodies, and protein loading is indicated by post-staining of Coomassie Brilliant Blue (CBB) of RuBisCO. This assay was conducted three times with similar results by RD. Cloning credits are provided in **Table S1**.

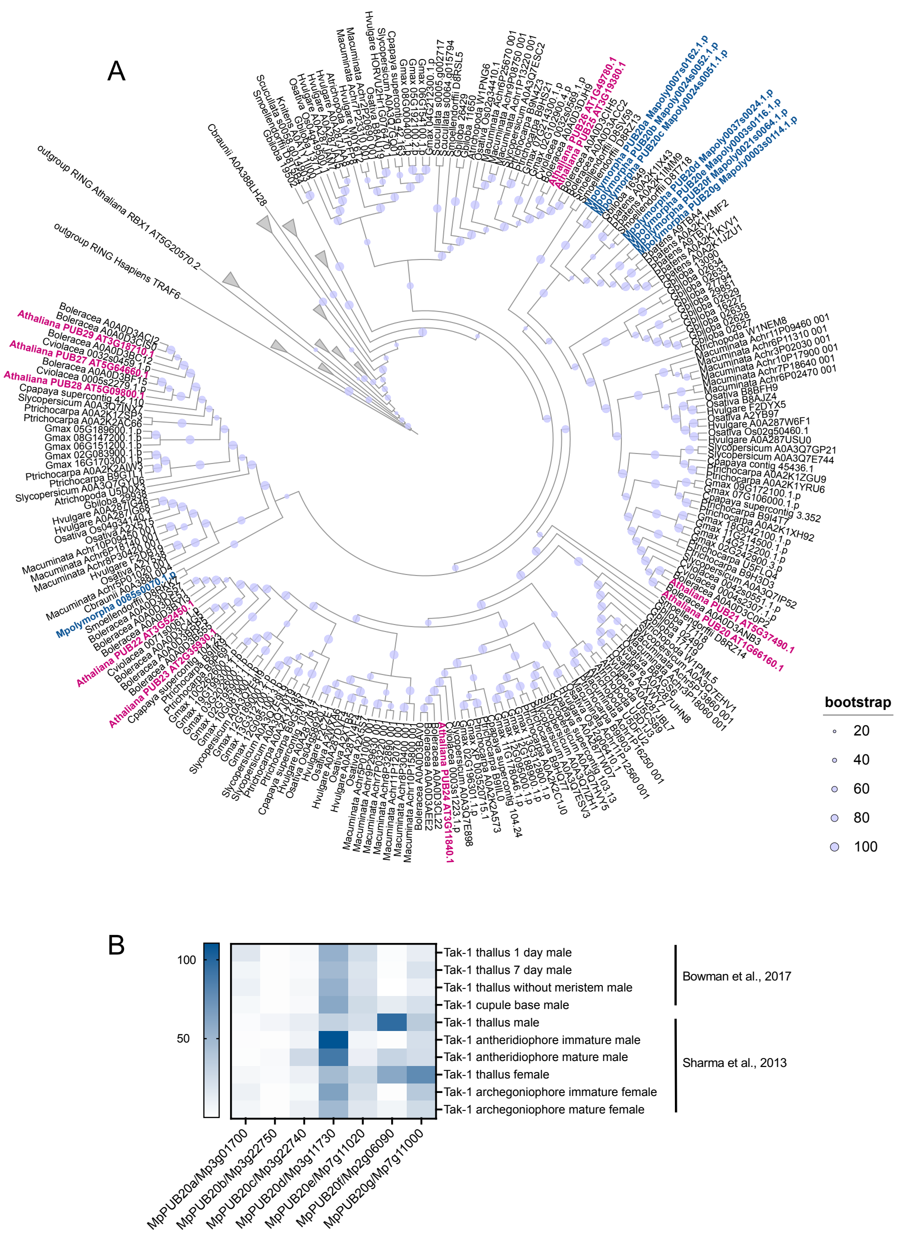


**Figure S5.** Identification of MpPUB20 genes in *Marchantia polymorpha*.

**(A)** Phylogenetic tree of a subset of class IV PUB protein sequences from 19 species across the plant lineage. The full phylogeny, including all details regarding its construction, is published as Figure S1 in (Trenner *et al.*, 2022); shown here is an excerpt of the relevant subfamily, visualized by JM in iTOL. Arabidopsis loci are coloured in magenta and Marchantia loci are coloured in blue. The gray circles indicate relative support of 2,000 bootstraps. **(B)** Heatmap showing expression data of MpPUB20 genes in the indicated tissues, based on data collected from two independent studies (Sharma *et al.*, 2013; Bowman *et al.*, 2022). Data was retrieved from MarpolBase and analyzed by JM.

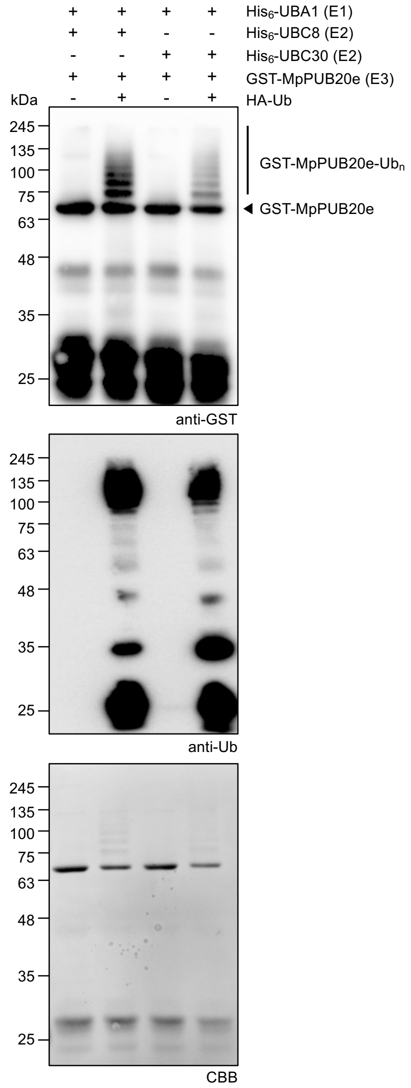


**Figure S6.** MpPUB20e is capable of auto-ubiquitination *in vitro* when paired with E2s AtUBC8 or AtUBC30.

*In vitro* ubiquitination assays using $His_6$-AtUBC1 as the E1 activating enzyme, $His_6$-AtUBC8 or $His_6$-AtUBC30 as the E2 conjugating enzyme, and GST-MpPUB20e as the E3 ligase enzyme. The upper western blot is probed with anti-GST; the middle blot is probed with anti-Ub; and protein loading is indicated by post-staining with Coomassie Brilliant Blue (CBB). These assays were repeated three times with similar results by RD. Cloning credits are provided in **Table S1**.

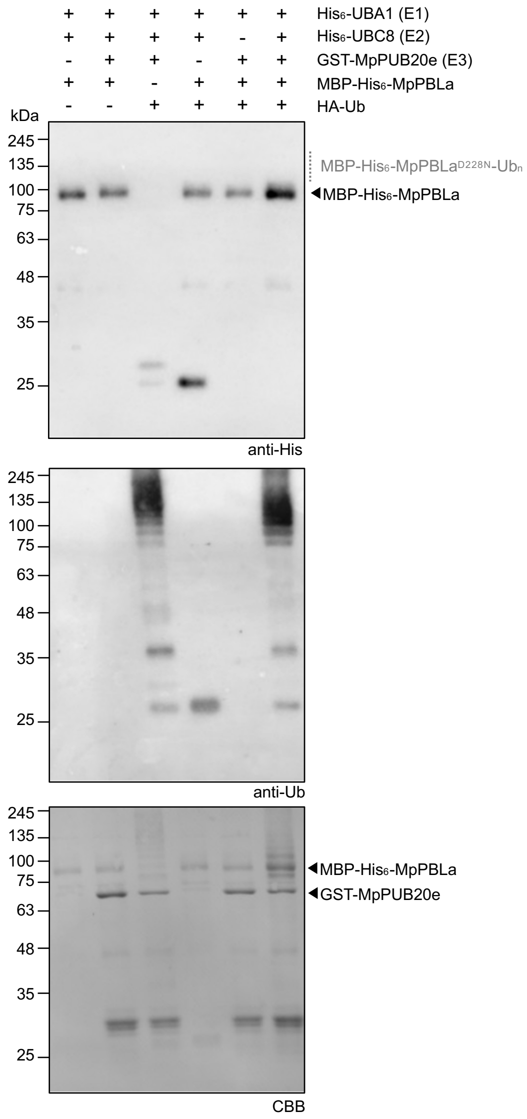


**Figure S7.** *In vitro* trans-ubiquitination of MpPBLa by MpPUB20e is not detectable.

*In vitro* ubiquitination assays using $His_6$-AtUBC1 as the E1 activating enzyme, $His_6$-AtUBC8 as the E2 conjugating enzyme, GST-MpPUB20e as the E3 ligase enzyme, and either MBP-$His_6$-MpPBLa as substrate. The upper western blots are probed with anti-His; the middle blot is probed with anti-Ub; and protein loading is indicated by post-staining with Coomassie Brilliant Blue (CBB). These assays were repeated more than three times with similar results by RD. Cloning credits are provided in **Table S1**.

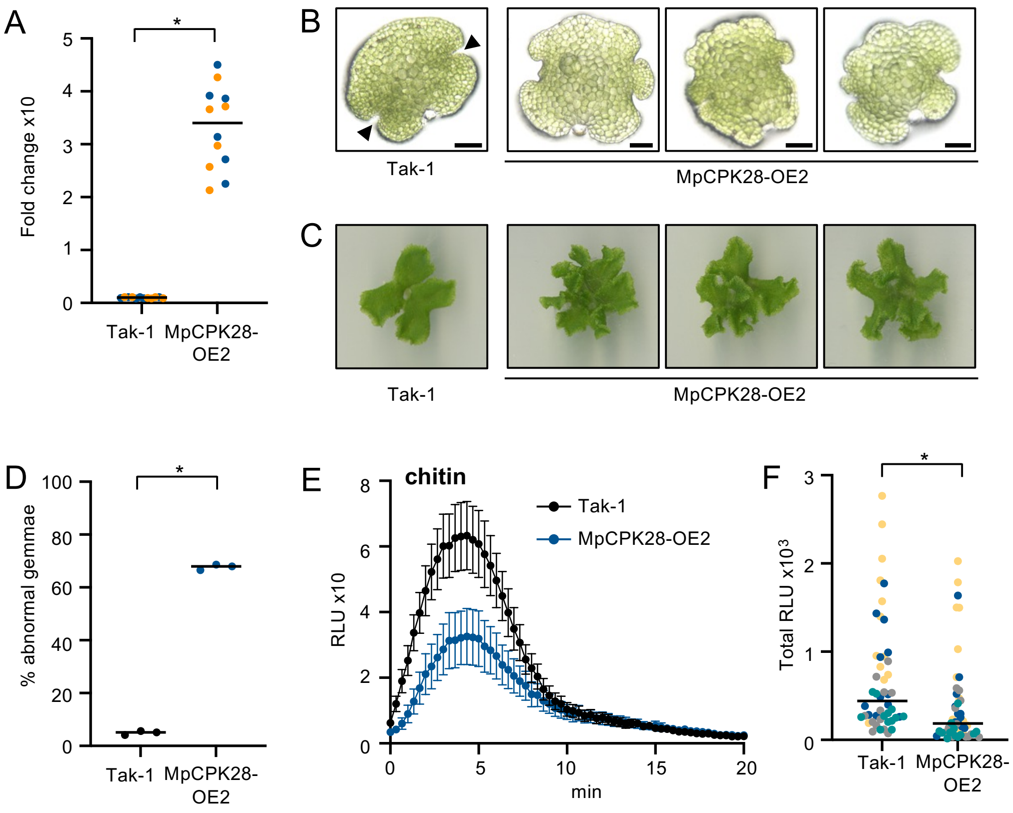


**Figure S8.** Characterization of MpCPK28-OE2 overexpression line in *Marchantia polymorpha*.

**(A)** Fold-change expression level of *MpCPK28* in the indicated genotypes relative to *MpACT* (blue dots) or *MpEF1a* (orange dots) in the indicated genotypes, as determined by quantitative reverse-transcription polymerase chain reaction (qRT-PCR). Values represent values from independent clones. Asterisk indicates a statistically significant difference according to an unpaired Student's t-test ($p$<0.0001). This experiment was repeated twice with similar results by DL. **(B)** Gemma from the indicated genotypes observed under a light microscope. Typical apical nodes are indicated by black arrowheads on Tak-1. Scale bar: 100 $\mu$m. **(C)** Representative thalli from 1-month-old plants of the indicated genotypes. Similar morphology was observed over several years by KEM. **(D)** Percent of abnormal gemmae, defined as those with more than two apical nodes, observed in the indicated genotypes. Values are means (n=100-150 gemma) from three independent biological replicates conducted by KEM. Asterisk indicates a statistically significant difference according to an unpaired Student's t-test ($p$<0.0001). **(E-F)** ROS production measured in relative light units (RLUs) after treatment with 0.1 mg/mL chitin. Values in **E** represent means +/- standard error (n=12 thallus discs) over a 20-min time-course from one representative experiment; values in **F** represent total RLU observed for each genotype, colour-coded by experimental replicate (n=4 replicates of 12 thallus discs each). Asterisk indicates a statistically significant difference between the indicated genotypes (unpaired Student's t-test; $p$=0.0118). Assays were repeated four times by JC. Cloning credits are provided in **Table S1**.

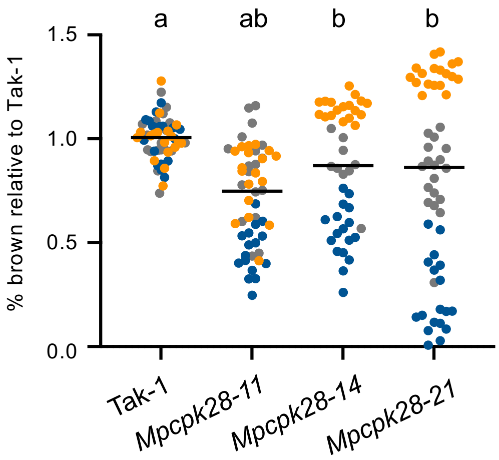


**Figure S9.** Infection of *Mpcpk28* mutants with *Collectotrichum nymphaeae.*

Three week old plants were drop-inoculated with 10 µL of a suspension of *Collectotrichum nymphaeae* spores (10,000 spores/mL water) and allowed to grow for another week. The ratio of thalli tissue that had turned brown after a week was assessed and plotted as % brown (diseased) compared to Tak-1 (raw values were based on measuring the diseased surface area in $cm^2$). Values represent individual infected plants colour-coded by experimental replicate and the line represents the mean across all technical replicates. Statistically significant groups are indicated by lower-case letters following a one-way analysis of variance with Tukey's posthoc test ($p$<0.0015).

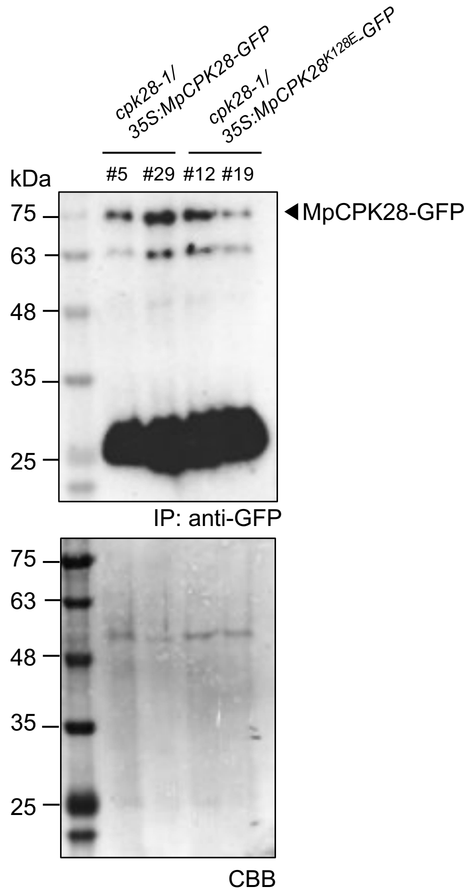


**Figure S10.** Western blot indicating MpCPK28-GFP expression in Arabidopsis transgenic lines.

Western blot indicating that MpCPK28-GFP and MpCPK28$^{K128E}$-GFP are expressed in the *cpk28-1* transgenic lines of the indicated genotypes. The proteins needed to be enriched by immunoprecipitation (IP) using GFP-Trap beads prior to western blotting with anti-GFP. Protein loading post-IP is indicated by post-staining with Coomassie Brilliant Blue (CBB) of RuBisCO. Similar protein expression patterns were observed over several replicates performed by MGD. Cloning and genotyping credits are provided in **Table S1**.

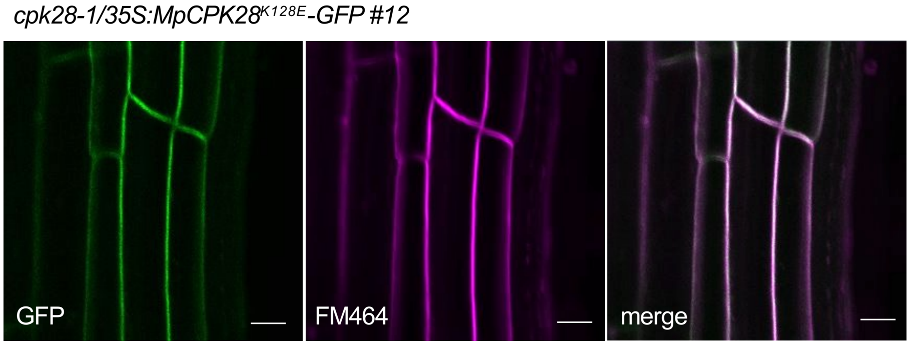


**Figure S11.** Subcellular localization of MpCPK28$^{K128E}$-GFP.

Single-plane confocal micrographs of root cells from *cpk28-1/35S:MpCPK28$^{K128E}$-GFP* line #12 indicating the subcellular localization of MpCPK28-GFP (GFP) and the plasma membrane (FM464). Scale bar indicates 10 µm. Data collected by RD.

### Supporting Information - Materials & Methods

#### Methods S1. Full details pertaining to the materials and methods used in this study.

##### Plant growth conditions

***Arabidopsis thaliana:*** *A. thaliana* plants (ecotype Col-0) were grown in controlled environment chambers (BioChambers and Conviron) in the Queen's University Phytotron. For soil assays, seeds were sown on Sunshine Mix 1 (Sungro) and seedlings transplanted to larger pots about a week after germination. Plants were grown under short-day conditions (8h light; 150 $\mu$M photons $m^2$ $s^{-1}$) at 22°C, watered when needed and supplemented with 1g/L fertilizer (20N:20P:20K) every 2 weeks. Mite bags containing *Amblyseius swirskii* (Koppert) were added to each tray of plants bi-weekly to prevent pest infestations. At the time of vegetative-to-reproductive transition (~6 weeks), plants were transferred to a chamber under long-day conditions (14h light) to assess stem elongation. For plate assays, seeds were surface-sterilized with 40% bleach for 30 minutes, rinsed well with sterile water, stratified in the dark at 4°C for 2-5 days, and sown on petri plates containing 0.5X Murashige & Skoog (MS) media and 0.8% agar. To assess growth inhibition, 4-day-old seedlings were transferred to single wells of 48-well plates containing liquid 0.5X MS media supplemented with 1% sucrose and the indicated concentration of AtPep1. After 12 days, individual seedlings were blotted dry and weighed on an analytical scale as described in (Bredow *et al.*, 2019). Transgenic lines were generated by floral dip with *Agrobacterium tumefaciens* GV3101 according to (Clough & Bent, 1998). Bacterial pellets were resuspended in 5% sucrose and 0.01% Silguard as a surfactant to an $OD_{600}$ = 0.5 and applied to unopened buds of ~5-week-old *A. thaliana* plants grown since germination in long-day conditions. Primary stems were trimmed to encourage axillary bud formation. Inoculated plants were kept in the dark overnight and seeds collected several weeks later. T1 plants were selected on 0.5X MS media with 0.8% agar and 50 ug/mL kanamycin prior to soil transfer. Homozygous lines containing single inserts were used for phenotyping, as indicated from segregation analysis in the T2 and T3 generations.

***Marchantia polymorpha:*** *Marchantia polymorpha ssp ruderalis* ecotype Takaragaike-1 (Tak-1) was cultivated axenically from gemmae and grown under a long day photoperiod (16 h light at 22°C/8h dark) at 20°C with 60-80 µE light intensity on 0.5X strength Gamborg B5 medium (Sigma, G5396), pH 5.7, 1.4% Agar-Agar (Sigma, A7921). Gemmae and thalli growth and development was captured using an Axio Zoom.V16 (Zeiss), GT-200 scanner (Epson), and analyzed with ImageJ. Transformation was conducted as previously described (Rich *et al.*, 2021), with some modifications. Freshly collected gemmae from axenic *M. polymorpha* Tak-1 were grown in 0.5X strength Gamborg B5 medium for five days. For each transformation, ~30 gemmalings were ground in a sterile, 250 mL stainless steel bowl (Waring, USA) in 10 mL of M51C (Takenaka et al., 2000) for 30 seconds. The grinded material was then transferred to a 250-mL erlenmeyer flask containing 40 mL of M51C medium and kept shaking (180 rpm) for three days at 22°C 16h light/ 20°C 8h dark. To co-culture with *A. tumefaciens* GV3101 for transformation, 100 $\mu$L of a saturated culture (pellet resuspended in 1 mL of M51C) was supplemented with 100 $\mu$M acetosyringone and 500 mg/mL glutamine. After three days, the content of the flasks was transferred into 50 mL conical tubes. Plant tissues were washed by decantation four times with sterile water, and plated on selective media: 0.5X Gamborg B5 with 200 mg/L amoxycilin (Levmentin, Laboratoires Delbert, FR) and 10 mg/L hygromycin (Duchefa Biochimie, FR). Chitin-induced ROS burst assays were performed on 4 week-old thalli as described in (Chu *et al.*, 2023). In brief, gemmae were cultured for 5 days in liquid 0.5X Gamborg B5 medium supplemented with 0.1 % sucrose at 22°C 16h light/ 20°C 8h dark with shaking. Gemmae were washed with sterile water and incubated in 20 $\mu$M 8-amino-5-chloro-7-phenylpyrido [3,4-d] pyridazine-1,4-(2H,3H) (L-012) (Wako, Japan) for 2 hours at 22 °C under darkness. The same volume of mock or elicitor solution was then added to a final concentration of 1 $\mu$M chitin and 2.5 U horseradish peroxidase (Sigma). ROS production was measured with a CLARIOstar platereader (BMG Labtech).

#### Phylogenetics

To explore the evolutionary history of CDPKs, we used HMMER v3.4 and HMMscan (hmmer.org) employing the complete PFAM and TIGERFAM datasets to thoroughly examine protein sequences containing a protein kinase (PF00069, TIGR03903) from the genomes of *Selaginella moellendorffii* (GCF_000143415.4), *Physcomitrium patens* (GCF_000002425.4), *Chara braunii* (GCA_003427395.1), *Klebsormidium nitens* (GCA_000708835.1), *Spirogloea muscicola* (GCA_009602725.1), *Oryza sativa* (Japonica Group; GCF_001433935.1), *Amborella tricopoda* (GCF_000471905.2), *Picea abies* (V1.0b), and *Marchantia polymorpha* (V6.r1). So that we could include the CRK subfamily, we did not further refine this search with PF00036 but rather compared sequences with an e-value >1.1e-30 to CDPK/CRK orthologs from *Arabidopsis thaliana* (TAIR10) using BLASTp 2.12.0+ to confirm homology (e-value 1e-5 allowing up to 10 targets). Five apicomplexan CPKs, including *Toxoplasma gondii* TgCPK1 (ToxoDB ID 162.m00001), *Toxoplasma gondii* TgCPK3 (ToxoDB ID 541.m00134), *Plasmodium falciparum* PfCPK3 (PlasmoDB ID PFC0420w), *Cryptosporidium parvum* CpCPK1 (CryptoDB ID cgd3_920), and *Cryptosporidium parvum* CpCPK3 (CryptoDB ID cgd5_820) were included in the alignments and used as outgroups. In total, 157 sequences were aligned using MAFFT v7.490 (Katoh & Standley, 2013) with the L-INS-i strategy, and sequences were trimmed using TrimAl v1.4.rev15 (Capella-Gutiérrez *et al*., 2009), which was employed for automated alignment trimming and removing poorly aligned sequences. The phylogenetic tree was reconstructed using IQ-TREE v2.3.4 (Minh *et al*., 2020), where the best-fitting evolutionary model (Q.plant+R6) was determined using ModelFinder (Kalyaanamoorthy *et al*., 2017) as implemented in IQ-TREE. The tree's robustness was assessed by applying Jackknife resampling with 1000 replicates, in addition to SH-aLRT (Guindon *et al*., 2010) and UltraFast Bootstraps

(Hoang *et al.*, 2018) for branch support, also using 1000 replicates. The final maximum likelihood tree was visualized and annotated in iTOL v6 (Letunic & Bork, 2021).

To explore the deep evolutionary history of subgroup IV CDPKs, sequences were retrieved from a database containing 193 plant species covering the Viridiplantae diversity using the tBLASTn v2.11.0+ (Camacho *et al.*, 2009) with an e-value threshold of 1-e10. Homolog nucleotide sequences were then aligned using MAFFT v7.471 (Katoh & Standley, 2013) and the FFT-NS-2 strategy. Subsequent alignment was then trimmed using trimAl v1.4 (Capella-Gutiérrez *et al.*, 2009) to remove positions with more than 60% of gaps. Finally, the tree was reconstructed using FastTree v2.1.10 (Price *et al.*, 2009) with default parameters and the -gtr option. From this tree, subtree corresponding to orthologs of MpCPK28 was pruned, and sequences aligned and trimmed as described above. Maximum likelihood tree was reconstructed using IQ-TREE v2.0.3 (Minh *et al.*, 2020) where the best-fitting evolutionary model (GTR+F+R7) was determined using ModelFinder as implemented in IQ-TREE (Kalyaanamoorthy *et al.*, 2017) according to the Bayesian Information Criteria (BIC). Branch supports were estimated using 10,000 replicates of both SH-aLRT (Guindon *et al.*, 2010) and UltraFast Bootstraps (Hoang *et al.*, 2018). Trees were visualized and annotated in iTOL v5 (Letunic & Bork, 2021).

#### Molecular cloning

Detailed information about molecular cloning of all vectors used in this study can be found in **Table S1**. We used several molecular cloning methods including Golden Gate, Gateway, In-Fusion, Gibson Assembly, and digestion-ligation cloning. For Golden Gate cloning, coding sequences were domesticated, synthesized and cloned into pMS (GeneArt, Thermo Fisher Scientific), and assembled as described in (Rich *et al.*, 2021). For Gateway cloning, recombination of entry vectors into Gateway-compatible binary vectors was achieved using Gateway LR Clonase II (Invitrogen) according to the manufacturer's instructions. Custom synthetic DNA fragments (Twist BioSciences or GeneArt) or PCR amplicons generated using Q5 *Taq* polymerase (NEB) and gene-specific primers, were used for In-Fusion cloning with the In-Fusion Snap Assembly

Master Mix (Takara), or for Gibson Assembly using the Gibson Assembly Master Mix (NEB), or for digestion-ligation cloning using construct-specific endonucleases (NEB) and T4 DNA ligase (NEB), all according to the manufacturer's directions. Mutations were incorporated either though PCR-mediated site-directed mutagenesis as described in (Monaghan *et al.*, 2014), or at the gene synthesis stage. Plasmids were transfected into *Escherichia coli* Top10 cells for replication and isolated using standard miniprep kits. Successful assembly of all plasmids was verified using Sanger sequencing or whole-plasmid sequencing (Plasmidsaurus).

#### RNA isolation, cDNA synthesis, and RT-qPCR

Total RNA was extracted from ~100 mg of ground frozen tissue from 2-3 weeks old *M. polymorpha* thalli using the Direct-zol RNA MiniPrep Zymo Kit (Ozyme) following the manufacturer's instructions. 1 µg of total RNA was treated with DNAse RQ1 (Promega) and cDNA was synthesized using the M-MLV Reverse Transcriptase Kit (Promega), both according to the manufacturer's instructions. Quantitative real-time PCR (RT-qPCR) was performed using 4 µL of 5-fold diluted cDNA, 5 µL of LightCycler 480 SYBR Green I Master Mix (Roche) and 1 µL of each primer at 10 µM, using the CFX Opus 384 Real-Time PCR System (Bio-Rad). Primers used for qPCR have an efficiency between 1.96-2.01 and are provided in **Table S1**.

#### Recombinant protein purification, *in vitro* kinase assays, and *in vitro* E3 ligase assays

All proteins were expressed and purified from *E. coli* strain BL21 or BL21-λP cells as recently described (Gonçalves Dias *et al.*, 2024). $His_6$-tagged proteins were purified using nickel-nitrilotriacetic acid (HisPur™ Ni-NTA, Thermo Fisher Scientific), and GST-tagged proteins were purified using pre-hydrated glutathione agarose beads (Sigma Aldrich). $His_6$-tagged proteins were eluted by sequential washes with imidazole-containing buffers at increasing concentrations (10 mM, 25 mM, 50 mM, 250 mM, and 500 mM) and dialyzed overnight at 4°C in 2000 volumes of buffer containing 25 mM Tris-HCl (pH 7.5), 50 mM

NaCl, and 1 mM DTT. GST-tagged proteins were eluted by incubation at 4°C overnight in a buffer containing 50 mM Tris-HCl (pH 7.5), 10 mM reduced glutathione, and 5 mM DTT. All proteins were concentrated using Amicon Ultra-15 or Ultra-4 centrifugal filter units (30 or 50 kDa MWCO; MilliporeSigma), and concentrations were determined using the Bradford assay (Thermo Fisher). Proteins were either immediately used in assays or aliquoted, flash-frozen in liquid $N_2$ and stored at –80°C until further use.

*In vitro* kinase assays using pIMAGO (Tymora Analytical) were performed according to the manufacturer's instructions and as previously described (Bredow *et al.*, 2021). $Ca^{2+}$-dependent autophosphorylation assays were performed by incubating 5 µg of purified recombinant $His_6$-MpCPK28 in a 50 µL reaction containing 25 mM Tris-HCl (pH 7.5), 10 mM $MgCl_2$, 1 mM DTT, 100 µM ATP, and either 100 µM $CaCl_2$ or 10 mM EGTA. The reaction was carried out at 30°C for 10 minutes. Reactions were terminated in 5× Laemmli sample buffer (LSB) at 80°C for 10 minutes prior to detection using pIMAGO. To assess CaM-mediated inhibition, 5 µg of $His_6$-MpCPK28 was incubated in a buffer containing 25 mM Tris-HCl (pH 7.5), 10 mM $MgCl_2$, 1 mM DTT, and 1 mM PMSF, with or without 100 µM $CaCl_2$ or 100 µM ATP, and with or without 2 µM CaM as described previously (Bender *et al.*, 2017). The reactions were incubated at 30°C for 30 minutes prior to the addition of 100 µM ATP to allow the formation of the MpCPK28-CaM complex. After ATP addition, the reactions proceeded for 30 seconds. Reactions were terminated in 5× LSB at 80°C for 10 minutes prior to detection using pIMAGO. Transphosphorylation assays using γP32-ATP were performed as recently described (Gonçalves Dias *et al.*, 2024), using 2 µg of kinase and 4 µg of substrate in a buffer containing 50 mM Tris-HCl (pH 8.0), 25 mM $MgCl_2$, 500 µM $CaCl_2$, 5 mM DTT, 5 µM ATP, and 2-4 µCi γ-[P32]-ATP, and incubated at 30˚C for 10 minutes. Reactions were terminated in 5× LSB at 80°C for 10 minutes, separated by SDS-PAGE. The resulting gels were sandwiched between two sheets of transparency film, exposed to a storage phosphor screen (Molecular Dynamics), and visualized using a Typhoon 8600 Imager (Molecular Dynamics). Gels were post-stained with Coomassie Brilliant Blue (CBB) R-250 (MP Biomedicals) or SimplyBlue SafeStain (Invitrogen; CBB G-250) and scanned.

*In vitro* E3 ubiquitin ligase assays were performed by incubating 2 µg of recombinant GST-MpPUB20e or variants and 75 µg/mL of HA-tagged ubiquitin (R&D Systems) in a buffer containing 25 mM Tris-HCl (pH 7.4), 5 mM $MgCl_2$, 25 mM KCl, 0.33 mM DTT, 1.5 mM ATP, 4 ng/µL E1 (UBA1), and 9 ng/µL E2 (UBC8 or UBC30). For trans-ubiquitination assays, 2 µg of substrate was additionally added. Reactions were incubated at 30°C for 10 - 180 min, terminated in 4× LSB at 80°C for 3 minutes, and finally separated by SDS-PAGE followed by immunoblot.

#### Phosphoproteomics

Phosphoproteomics was conducted exactly as described in (Gonçalves Dias *et al.*, 2024) without deviation, with the acquired LC-MS data searched against the *Marchantia polymorpha* v3.1 proteome available on Phytozome (https://phytozome-next.jgi.doe.gov/). Proteomics data has been deposited to the ProteomeXchange Consortium via the PRIDE (https://www.ebi.ac.uk/pride/) partner repository with the data set identifier PXD056365. Username:; Password: w9uEckCrRqPJ.

#### Cell-free degradation assays

Tissue from Marchantia Tak-1 (3- to 4-week-old plants grown on ½ B5 medium) was collected and ground to a fine powder in liquid $N_2$. Total protein was extracted in a buffer containing 50 mM Tris-MES (pH 8.0), 0.5 M sucrose, 10 mM EDTA (pH 7.5 or 8.0), 5 mM DTT, and 1 mM $MgCl_2$ at a ratio of approximately 1 mL buffer per ~1 mL ground tissue. The mixture was incubated at 4°C with shaking for 1-1.5 hours, followed by centrifugation at 14,000 × *g* for 30 minutes at 4°C. The supernatant was filtered through miracloth to remove any remaining plant matter. Protein concentration was determined using a Bradford assay (Thermo Fisher), and the extract was normalized to 1.0 mg/mL. For the degradation assay, 1.0 mg/mL of freshly extracted protein was mixed with 1 µL of 1.5 mg/mL recombinant proteins and 10 mM ATP in a 400 µL reaction incubated at 30°C. If MG132 proteasome inhibitor was used, a 50 $\mu$L sample was taken at 0 hour, mixed with

0.5 *μ*L of 10 mM MG132, and the reaction was stopped at 6 hours. A 50 µL sample was taken every 2 hours, and the reaction was stopped by adding 5× LSB at 80°C for 10 minutes. The samples were subjected to SDS-PAGE followed by immunoblotting.

#### Immunoblots

Proteins were separated by 6-10% SDS-PAGE, transferred onto polyvinylidene difluoride membranes (Bio-Rad), and blocked with 5% nonfat milk or bovine serum albumin (BSA) in Tris-buffered saline containing 0.05–0.1% Tween-20. All antibodies used are listed in **Table S1**. Membranes were incubated with ECL Clarity Substrate (Bio-Rad), visualized on a ChemiDoc Touch Imaging System (Bio-Rad) and subsequently stained with Coomassie Brilliant Blue (CBB) R-250 (MP Biomedicals) as a loading control. Detection of MpCPK28-GFP in *cpk28-1* transgenic lines was facilitated by immunoprecipitation using GFP-Trap beads as described previously (Monaghan *et al*., 2014).

#### Confocal microscopy

Images were taken using a Zeiss LSM710 confocal microscope in the Biology Department at Queen's University and processed using Zeiss Zen Software. Roots of Arabidopsis seedlings were stained with 2 µM FM464 (Calbiochem) prior to mounting in water on a glass slide. Fluorescent proteins were excited with a 488 nm Argon laser and imaged using separate channels to detect emission of GFP (510-540 nm) or RFP (635-680 nm).

#### Pathogen infection assays

Arabidopsis plants were infected with *Pseudomonas syringae* pathovar *tomato* isolate DC3000 on 4-5 week old plants grown in short-day conditions exactly as previously described (Monaghan *et al.*, 2014). Marchantia plants were infected with *Pseudomonas viridiflava* isolate MPG032 as described previously (Robinson *et al.*, 2025). Briefly, 3-week old plants grown on G1/2 media were vacuum infiltrated with a solution of

*Pseudomonas viridiflava* MPG032 at $OD_{600}$ = 0.02 in 10 mM $MgCl_2$ and bacteria were isolated and plated by serial dilution 3 days post infection. Marchantia plants were infected with *Collectotrichum nymphaeae* as described previously (El Mahboubi *et al.*, 2026). Briefly, 3-week-old plants grown on G1/2 media were drop inoculated with 10 µL of a solution of 10,000 *Colletotrichum nymphaeae* spores per mL water solution. Plants were photographed one week after inoculation and the ratio of brown areas in $cm^2$ (indicative of disease) to green areas (indicative of resistance) was measured by hand using ImageJ.

#### Quantification and Statistical Analyses

Data were graphed using GraphPad Prism v10. Parameters for all statistical tests are described in the figure legends.

#### Accession numbers

Mp4g01600/Mapoly0098s0040//MpCPK28;

Mp3g25360/Mapoly0100s0049/MpPBLa;

Mp3g01700/Mapoly0007s0162/MpPUB20a;

Mp3g22750/Mapoly0024s0052/MpPUB20b;

Mp3g22740/Mapoly0024s0051/MpPUB20c;

Mp3g11730/Mapoly0037s0024/MpPUB20d;

Mp7g11020/Mapoly0003s0116/MpPUB20e;

Mp2g06090/Mapoly0021s0064/MpPUB20f;

Mp7g11000/Mapoly0003s0114/MpPUB20g.

### Supporting Information - References

**Bender KW, Blackburn RK, Monaghan J, Derbyshire P, Menke FLH, Zipfel C, Goshe MB, Zielinski RE, Huber SC**. **2017**. Autophosphorylation-based Calcium (Ca2+) Sensitivity Priming and Ca2+/Calmodulin Inhibition of Arabidopsis thaliana Ca2+-dependent Protein Kinase 28 (CPK28). *The Journal of Biological Chemistry* **292**: 3988–4002.

**Bowman JL, Arteaga-Vazquez M, Berger F, Briginshaw LN, Carella P, Aguilar-Cruz A, Davies KM, Dierschke T, Dolan L, Dorantes-Acosta AE, *et al.* 2022**. The renaissance and enlightenment of Marchantia as a model system. *The Plant Cell* **34**: 3512–3542.

**Bredow M, Bender KW, Johnson Dingee A, Holmes DR, Thomson A, Ciren D, Tanney CAS, Dunning KE, Trujillo M, Huber SC, *et al.* 2021**. Phosphorylation-dependent subfunctionalization of the calcium-dependent protein kinase CPK28. *Proceedings of the National Academy of Sciences of the United States of America* **118**:e2024272118.

**Bredow M, Sementchoukova I, Siegel K, Monaghan J**. **2019**. Pattern-Triggered Oxidative Burst and Seedling Growth Inhibition Assays in Arabidopsis thaliana. *Journal of Visualized Experiments.* Doi: 10.3791/59437.

**Camacho C, Coulouris G, Avagyan V, Ma N, Papadopoulos J, Bealer K, Madden TL**. **2009**. BLAST+: architecture and applications. *BMC Bioinformatics* **10**: 421.

**Capella-Gutiérrez S, Silla-Martínez JM, Gabaldón T**. **2009**. trimAl: a tool for automated alignment trimming in large-scale phylogenetic analyses. *Bioinformatics* **25**: 1972–1973.

**Chu J, Monte I, DeFalco TA, Köster P, Derbyshire P, Menke FLH, Zipfel C**. **2023**. Conservation of the PBL-RBOH immune module in land plants. *Current Biology* **33**: 1130–1137.e5.

**Clough SJ, Bent AF**. **1998**. Floral dip: a simplified method for Agrobacterium-mediated transformation of Arabidopsis thaliana. *The Plant Journal* **16**: 735–743.

**El Mahboubi K, Beaulieu C, Castel B, Libourel C, Jariais N, Amblard E, van Beveren F, Keller J, Martinez Y, Nelson JM, *et al.* 2026**. Plant-fungi interactions in Marchantia polymorpha are associated with horizontal gene transfer and terpene metabolism. *Proceedings of the National Academy of Sciences of the United States of America* **123**: e2532723123.

**Gonçalves Dias M, Doss B, Rawat A, Siegel KR, Mahathanthrige T, Sklenar J, Rodriguez Gallo MC, Derbyshire P, Dharmasena T, Cameron E, *et al.* 2024**. Subfamily C7 Raf-like kinases MRK1, RAF26, and RAF39 regulate immune homeostasis and stomatal opening in Arabidopsis thaliana. *The New Phytologist* **244**: 2278–2294.

**Guindon S, Dufayard J-F, Lefort V, Anisimova M, Hordijk W, Gascuel O**. **2010**. New algorithms and methods to estimate maximum-likelihood phylogenies: assessing the performance of PhyML 3.0. *Systematic Biology* **59**: 307–321.

**Hoang DT, Chernomor O, von Haeseler A, Minh BQ, Vinh LS**. **2018**. UFBoot2: Improving the

Ultrafast Bootstrap Approximation. *Molecular Biology and Evolution* **35**: 518–522.

**Kalyaanamoorthy S, Minh BQ, Wong TKF, von Haeseler A, Jermiin LS**. **2017**. ModelFinder: fast model selection for accurate phylogenetic estimates. *Nature Methods* **14**: 587–589.

**Katoh K, Standley DM**. **2013**. MAFFT multiple sequence alignment software version 7: improvements in performance and usability. *Molecular Biology and Evolution* **30**: 772–780.

**Letunic I, Bork P**. **2021**. Interactive Tree Of Life (iTOL) v5: an online tool for phylogenetic tree display and annotation. *Nucleic Acids Research* **49**: W293–W296.

**Minh BQ, Schmidt HA, Chernomor O, Schrempf D, Woodhams MD, von Haeseler A, Lanfear R**. **2020**. IQ-TREE 2: New Models and Efficient Methods for Phylogenetic Inference in the Genomic Era. *Molecular Biology and Evolution* **37**: 1530–1534.

**Monaghan J, Matschi S, Shorinola O, Rovenich H, Matei A, Segonzac C, Malinovsky FG, Rathjen JP, MacLean D, Romeis T, *et al.* 2014**. The calcium-dependent protein kinase CPK28 buffers plant immunity and regulates BIK1 turnover. *Cell Host & Microbe* **16**: 605–615.

**Price MN, Dehal PS, Arkin AP**. **2009**. FastTree: computing large minimum evolution trees with profiles instead of a distance matrix. *Molecular Biology and Evolution* **26**: 1641–1650.

**Rich MK, Vigneron N, Libourel C, Keller J, Xue L, Hajheidari M, Radhakrishnan GV, Le Ru A, Diop SI, Potente G, *et al.* 2021**. Lipid exchanges drove the evolution of mutualism during plant terrestrialization. *Science* **372**: 864–868.

**Robinson K, Buric L, Grenz K, Chia K-S, Hulin MT, Ma W, Carella P**. **2025**. Conserved effectors underpin the virulence of liverwort-isolated Pseudomonas in divergent plants. *Current Biology* **35**(8):1861-1869.e3.

**Sharma N, Bhalla PL, Singh MB**. **2013**. Transcriptome-wide profiling and expression analysis of transcription factor families in a liverwort, Marchantia polymorpha. *BMC Genomics* **14**: 915.

**Trenner J, Monaghan J, Saeed B, Quint M, Shabek N, Trujillo M**. **2022**. Evolution and Functions of Plant U-Box Proteins: From Protein Quality Control to Signaling. *Annual Review of Plant Biology*. **73:** 93-121.
